# Microchannel patterning strategies for *in vitro* structural connectivity modulation of neural networks

**DOI:** 10.1101/2021.03.05.434080

**Authors:** B. G. C. Maisonneuve, J. Vieira, F. Larramendy, T. Honegger

**Affiliations:** Univ. Grenoble Alpes, CNRS, LTM, 38000 Grenoble, France; NETRI, 69007 Lyon, France

## Abstract

Compartmentalized microfluidic chips have demonstrated tremendous potential to create *in vitro* minimalistic environments for the reproduction of the neural circuitry of the brain. Although the protocol for seeding neural soma in these devices is well known and has been widely used in myriad studies, the accurate control of the number of neurites passing through the microchannels remains challenging. However, the regulation of axonal density among different groups of neurons is still a requirement to assess the inherent structural connectivity between neuronal populations. In this work, we report the effect of microchannel patterning strategies on the modulation of neuronal connectivity by applying dimensional modifications on microchannel-connected microfluidic chambers. Our results show that those strategies can modulate the direction and the number of neuronal projections of passage, therefore regulating the strength of the structural connections between two populations of neurons. With this approach, we provide innovative microfluidic design rules for the engineering of *in vitro* physiologically relevant neural networks.

## Introduction

The reconstruction of neural circuits through microfluidics^1^ appears to be an efficient approach to build *in vitro* compartmentalized neural networks compared to conventional tissue culture techniques using petri dishes^4,5^. Neuronal connections are indeed built randomly based on cell body proximity, or brain slices-on-a-chip^2^, where connections create intermixed networks that are partially damaged as a cause of the slicing process of the tissue. The main advantage of microfluidic-based technologies is their ability to isolate and manipulate the cellular environment^3^, enabling the fabrication of simplistic, yet relevant, neural networks between different co-cultured neuronal cell types^4^. Translating the architecture of neural networks to *in vitro* models requires the rationalization of microfluidic design rules by regulating the connectivity among different neuronal populations. This connectivity has been already implemented through introducing axonal guidance systems in microfluidic chips that apply directionality to the developing neurites. Examples of these methods are the addition of microgrooves between microchannels^1^, the use of triangular^5^ and rerouting^6^ axonal diodes, increase of microchannel number^7^, and electrokinetic confinement^8,9^. However, to fully reconstruct relevant neural architectures *in vitro* it is essential to control not only the axonal directionality, but also the transmission weight of the connections, which are partly determined by the number of axons innervating dendrites. Both features are particularly important to understand the mechanisms of several currently incurable neurodegenerative diseases, such as Parkinson’s disease (PD). For example, an alteration of the nigrostrial path is firstly involved in the basal ganglia loop in PD, leading to deleterious dendritic remodelling and disrupted synaptic communication^10,11^. Therefore, there is a need to explore microfluidic strategies that can accurately reproduce such connectivity configurations *in vitro* to fully comprehend the nature of PD and other disorders, and advance in our ability to treat them. Here, we investigate whether variations in microchannel width and number can influence changes on the quantity of axons that project from one chamber to the other. Our results show that those adjustments can indeed be used to modulate the density of projected axons among compartmentalized neuronal clusters, suggesting that microchannel patterning strategies offer enhanced control over structural connectivity between neuronal populations.

## Materials and methods

### Device design and masks fabrication

The fabricated microfluidic device is a two-layered compartmentalized chip with two separated culture chambers connected by microchannels (Figure 1A). The design of the bottom layer (3-μm thick) contains microchannels of 500-μm long that are grouped in sets of widths ranging from 5 to 100 μm, where supporting pillars avoid the collapse of microchannels for widths greater than 50 μm (inset Figure 1A). The microchannels are narrow enough to exclusively allow the passage of neuronal projections to the second microfluidic chamber, while neuronal bodies remain within the first one (Figure 1B), and long enough to avoid that neurites from glial cells, such as astrocytes, are considered in the analysis^12^. The upper layer presents identical geometry for both chambers but does not include the microchannels patterns. This mask is used to increase the final height of the loading chambers to 100 μm.

**Figure 1:**
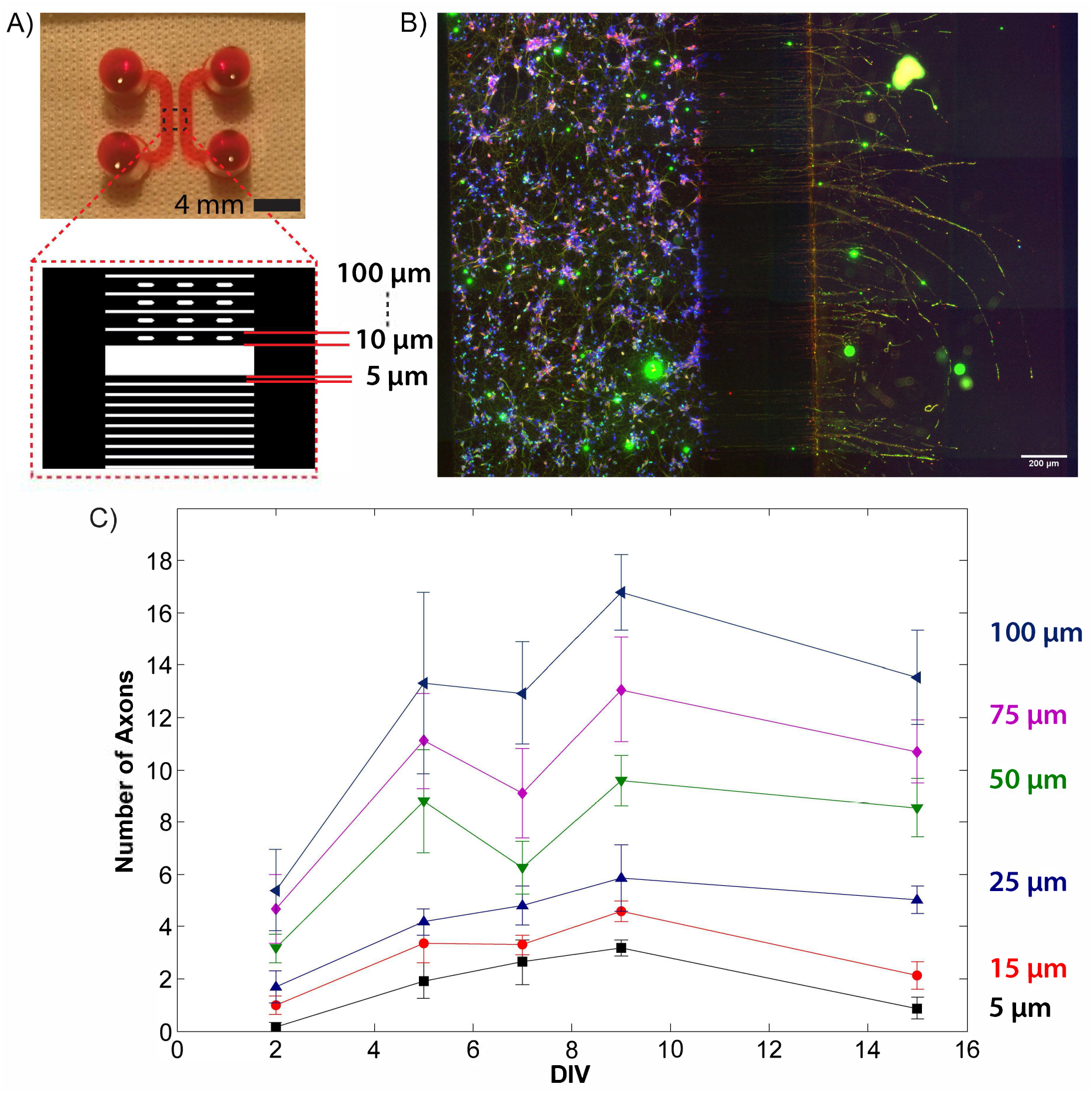
***(A)*** Image showing the general structure of the microfluidic device filled with rhodamine B. *Inset:* Schematic representation of the different widths of the microchannels. ***(B)*** Rat hippocampal neurons were fixed at DIV 21 and stained against Tuj1 (green) and Map2 (red). Neuronal nuclei were stained using DAPI (blue). The image was obtained using a 10x objective. ***(C)*** Quantification of number of projected axons through the microchannels of various widths as a function of days *in vitro* (DIV) when no extra volume of media was added (0 μL hydrostatic condition). Data indicated as mean ± SEM; (n = 3).

Respective masks for the designed layers were fabricated using a homemade mask-less UV photolithography system^13^, or purchased on transparent films (Selba SA, CH).

### SU-8 mold fabrication

This procedure was adapted from one of our previously published protocols^13^. The mold was fabricated using a silicon wafer as substrate. An adhesion promoter (Omnicoat, MicroChem, USA) was spin-coated onto the wafer at 300 rpm for 30 sec, before letting it bake at 200 °C for 1 min. SU8-2002 (MicroChem, USA) was spin-coated onto the wafer at 1000 rpm for 30 sec, followed by a soft bake step (1 min at 65 °C, 2 min at 95 °C, and 1 min at 65 °C). Using the mask corresponding to the bottom layer of the microfluidic chip, the wafer was exposed to an appropriate dose of UV light (90 mJ/cm²) using a MJB4 mask aligner (Süss MicroTech, Germany). After exposure, another soft bake step was done (1 min at 65 °C, 2 min at 95 °C, and 1 min at 65 °C) before developing the wafer in SU8 developer (MicroChem, USA). The wafer was finally rinsed with isopropanol. SU8-2050 was then spin-coated onto the same wafer at 1667 rpm for 30 sec, followed by another soft bake step (5 min at 65 °C, 16 min at 95 °C, and 5 min at 65 °C). The second mask was aligned with the SU8 features already onto the wafer using alignment marks, before exposing the wafer to UV light (230 mJ/cm²). A post-exposure bake step was performed (4 min at 65 °C, 9 min at 95 °C, and 2 min at 65 °C) and the wafer was developed in SU8 developer again, before being washed with isopropanol. A final hard bake step (150 °C for 15 min) was done.

### PDMS microfluidic chip fabrication

This procedure was adapted from one of our previously published protocols^13^. The wafer was first silanized using vapours of a silanizing agent (trichloro(1H,1H,2H,2H-perfluorooctyl) silane) in a desiccator for 30 min. Polydimethylsiloxane (PDMS) prepolymer (Sylgard 184, Dow Corning, USA) was prepared by mixing the base and the curing agent at a ratio of 10:1. The mixture was degassed, poured onto the wafer and cured at 80 °C for 40 min. The PDMS layer was cut to the required size, peeled off the wafer, and the inlet/outlet zones were punched out. The device and a clean microscope glass slide were then plasma treated using a plasma cleaner (Harrick Plasma, USA) before their assembly. The devices were then sprayed and filled with a solution of 70% ethanol and brought into a sterile environment. The ethanol was washed away three times using sterile distilled water and exposed to UV light for 30 min.

The channels within the chambers of the microfluidic device were then coated with 0.1 mg/mL poly-L-lysine (Sigma Aldrich, USA) and incubated for 24 hours at 37 °C. The channels were then rinsed three times with Hank’s Balanced Salt Solution (HBSS) (Life Technology, Thermo Fisher Scientific Inc., USA) buffered with 10mM 4-(2-hydroxyethyl)-1-piperazineethanesulfonic acid (HEPES) 1M (ThermoFisher, ref. 15630106) and coated with 20 μg/mL laminin (Sigma Aldrich, USA) for 2 hours. The coated channels were washed again with HBSS and filled three times with hippocampal culture medium composed of Neurobasal-B27 (ThermoFisher, ref. 17504044) with a mix of Penicillin-Streptomycin-Glutamine (ThermoFisher, ref. 10378016) containing 2 mM glutamine and 100 U/mL penicillin/streptomycin. The microfluidic chips were finally placed in an incubator until use.

### Neuron preparation and seeding

According to previously published procedures^13^, hippocampi were harvested from E18 OFA rats (Charles River Laboratories) and kept in ice-cold HBSS buffered with 10mM HEPES (pH 7.3). The tissue was digested for 30 min using 2 mL of HEPES-buffered HBSS containing 20 U/mL of papain (Worthington Biochem.), 1 mM EDTA (PanReac AppliCham, ref. A4892,0100) and 1 mM L-cysteine (SigmaAldrich, ref. 52-90-4 Then, the tissue was rinsed three times with 8 mL of hippocampal culture medium. The cells were gently triturated in 1 mL of hippocampal culture medium, counted with an hemocytometer, and seeded into the device. The cells were maintained under incubation conditions (37 °C, 5% CO2, and 80% humidity) until use.

Before seeding, the reservoirs of the microfluidic chip were emptied without removing the media from the channels. Through one inlet reservoir, 4 μL of high density (> 8106 cells/mL) dissociated neuron solution were placed near the entrance of the channel. The chip was replaced to the incubator for 5 min in order to let the neurons adhere on the coated surface, and the seeding process was repeated three times to achieve a high cell density. Finally, each input and output reservoirs of the device were filled with hippocampal culture medium, and chips were returned to the incubator.

All animal work performed in this project was approved by the CEA and CNRS Ethics Committee of Animal Care and abided by institutional and national guidelines for animal welfare.

### Neuron culture in device

The neurons seeded within the device were cultured up to 21 days *in vitro* (DIV), and the culture media was renewed up to three times per week. Cultured cells were subjected to three hydrostatic conditions, where the difference of static pressure at the air/fluid interface was controlled between the two microfluidic chambers through the addition of extra medium at the inlet and outlet of the culture chamber: (i) no addition of extra volume of medium (0 μL), (ii) addition of 40 μL of extra medium, and (iii) addition of 80 μL of extra medium. The difference in volume induces a height variation of the free surface and, therefore, a difference in hydrostatic pressure. Taking into consideration that the inlet and outlet cylindrical openings of the chambers have a radius of 2mm, the addition of 40 μL and 80 μL simulate a pressure increase of 30 Pa and 60 Pa, respectively. The induced hydrostatic pressure was calculated from the applied increase of volume by using Bernoulli’s principle.

### Cell number quantification

To quantify the number of cells that were present within the chambers, DIV 21 neurons were fixed with a solution of 4% paraformaldehyde (PFA) (Sigma-Aldrich) diluted in 1XPBS (Sigma-Aldrich) for 20 min. Next, neurons were rinsed five times with 1XPBS, and posteriorly permeabilized and blocked against nonspecific binding using a solution of 3% bovine serum albumin (BSA) (Sigma-Aldrich) and 0.1% Triton X-100 (Sigma-Aldrich) diluted in 1XPBS for 45 min. Neurons were subsequently washed five times with 1XPBS and incubated with a 300 nM DAPI solution (Thermo Fisher) for 10 min. After incubation, cells were washed five times with 1XPBS. Image acquisition was done using an epifluorescence microscope (CKX41, Olympus), and neuronal quantification was performed with a homemade MATLAB script (MATLAB 2019b, The MathWorks, Inc., USA).

### Immunostaining

The number of neurons in each device was marked using DAPI staining. The neurons culture were fixed with a 4% paraformaldehyde (PFA) (Sigma-Aldrich) solution in PBS (Sigma-Aldrich) for 20 minutes. The cells were rinsed 5 times with PBS, and the cells were permeabilized and blocked with a 3% bovine serum albumin (BSA) (Sigma-Aldrich) 0.1% Triton X-100 (Sigma-Aldrich) solution in PBS for 45 minutes. The cells were then rinsed again 5 times with PBS and a 300 nM DAPI solution (Thermo Fisher) was added for 10 minutes.

### Axonal number quantification

For axonal quantification analysis, neurons in culture were imaged at several time points: DIV 2, 5, 7, 9, and 15. For each time point, neurons were fixed and stained with DAPI as described in the previous section. Before imaging, neuronal bodies and projections were labelled with antibodies against MAP2 (Thermo Fisher, dilution 1:1000, secondary antibody: goat anti-rabbit Alexa Fluor 647, Life Technologies, dilution 1:1000) and Tuj1 (Abcam, dilution 1:1000, secondary antibody: goat anti-chicken Alexa Fluor 488, Life Technologies, dilution 1:1000). Image acquisition was done using an epifluorescence microscope (CKX41, Olympus) with 10x and 20x objectives, and the obtained images were stitched with Image Composite Editor (ICE, Microsoft) to reconstruct the entire device. The reconstructed images were used to manually count the number of axons that were exiting the 500-μm long microchannels to the chamber where no neurons were seeded.

### Statistical analysis

Statistical analysis was determined using one-way ANOVA with a post-hoc Tukey’s test. The statistical software used for the analysis was Origin Pro 8.0 (OriginLab Corporation, USA). All experiments were independently repeated at least three times. P-values and significancy are defined in the figure legends. On the figures, the extremities of the bars indicate which two conditions are compared, and the braces designate the comparison of more than two conditions.

## Results and discussion

### Influence of microchannel width variations on the number of projected axons

To investigate the influence of the variable width of the microchannels on the number of projected neurites, we designed a chip where the width differed between microchannels (Figure 1A). We evaluated the filtering abilities of the variations applied in our design by only seeding rat hippocampal neurons on one side of the device, similarly to previous protocols for design characterization^5,6^. Seeded neurons were maintained on culture until DIV 21 and the number of projected axons was evaluated at different time points by immunostaining against MAP2 and TUJ1 (Figure 1B). Although these particular proteins used are not specific for axons, previous studies have shown that dendrites are found only in the proximal part of the guiding structures, and that almost all the traversing neuronal projections that exit microchannels longer than 450 μm can be considered axons^1,7,14^.

Results show that, already at DIV 2, axons start to exit the microchannels (Figure 1C). The number of projected axons increases with time until neurons reach DIV 9, and this amount does not change significantly between DIV 9 and DIV 15 (Figure 1C). This finding confirms that, the wider the microchannels are, the more neurites penetrate trough them, which is consistent with the natural projection of neurites from one chamber to the other^1,5,6^. We also identified numerous axons that presented a non-linear shape within microchannels more than 10-μm wide (data not shown), indicating an alternative growing path that does not follow the orientation of the microchannels. This observation is consistent with previous studies, where growth cone exploration during axonal expansion within microchannel confinement was identified^5,15^.

Based on the fact that an increasing number of axons was detected between DIV 2 and DIV 9, we decided to use these two culture time points in our experiments as comparable conditions, and thus to be able to observe substantial differences on axonal growth and density.

### Influence of neuronal density on the number of developed axons

Next, we questioned whether the variations on the number of projected axons along microchannels could be partly affected by differences on neuronal density between experiments. The maintenance of a uniform distribution of seeded cells within the microfluidic chamber across experimental repetitions it is extremely challenging, even following the same seeding procedure for every technical replicate. In order to control for this variability in our data, the neurons seeded on the coated channels were stained with DAPI at DIV 9 to perform nuclei quantification after imaging. As expected, results indicate a highly heterogenous number of counted cells in relation to their position within the microfluidic channel (Figure 2A).

**Figure 2:**
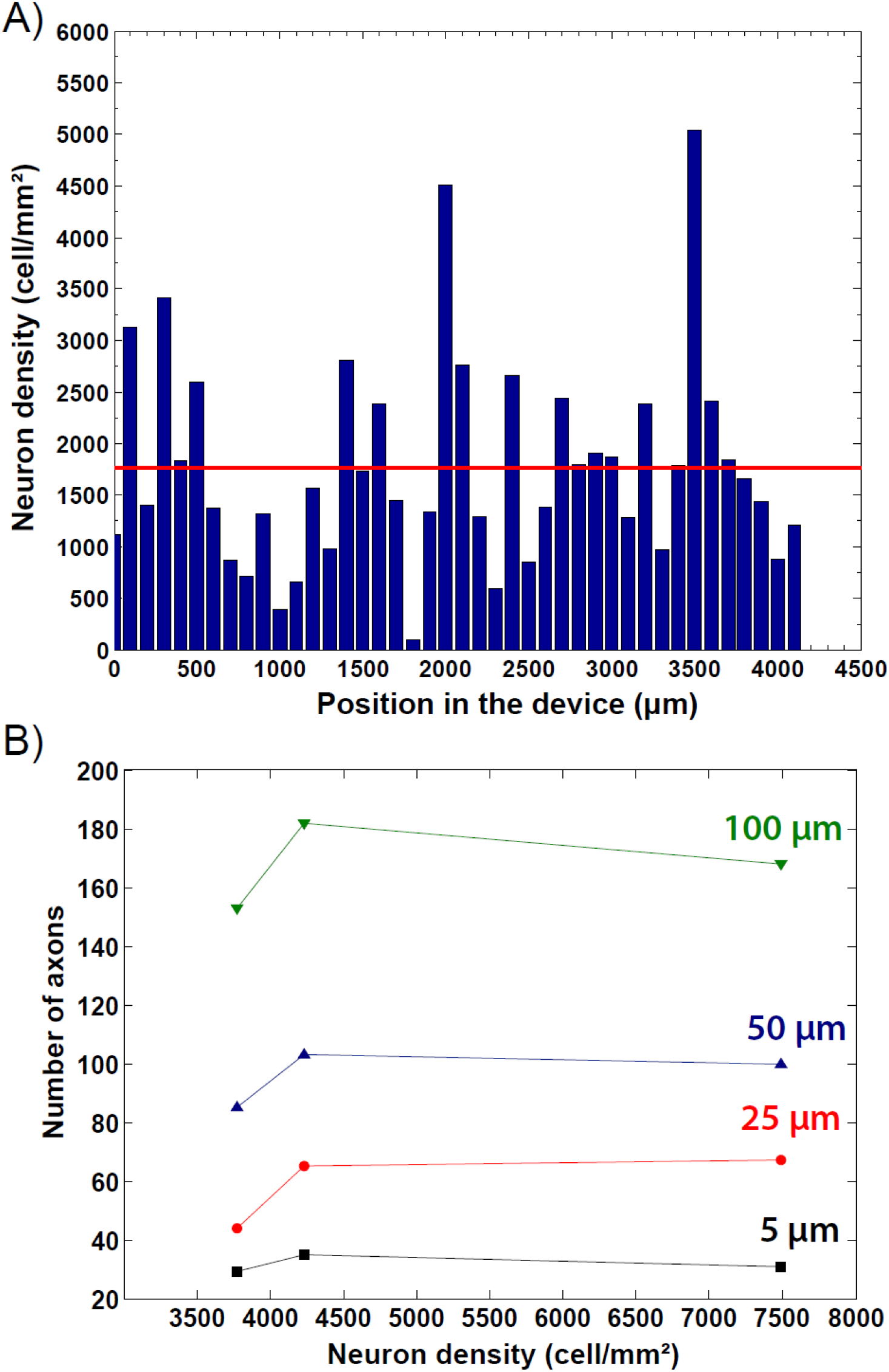
***(A)*** Density of seeded neurons (cells/mm^2^) as a function of their position along the culture chamber of the microfluidic device. The red line shows the mean value of approximately 1750 neurons/mm². ***(B)*** Quantification of number of projected axons exiting the microchannels of different widths as a function of the density of seeded neurons (cells/mm^2^) at DIV 9 when no extra volume of media was added (0 μL hydrostatic condition). Data indicated as mean; (n = 3).

To further investigate the impact of those neuronal density irregularities on the number of projected axons, the quantified number of axons going through microchannels of 5, 25, 50, and 100 μm width was divided by the different cell densities tested within the culture chambers. Importantly, our data show that the number of counted axons does not significantly vary in relation to the neuronal density (Figure 2B), suggesting that unequal neuronal distribution along the seeding chamber is not influencing the resulting axonal density.

### Normalization regarding geometrical constraints

Knowing that the wider the microchannel in our device, the more axons of passage (Figure 1A), we next aimed to compare the average number of projected axons counted within a single 100-μm wide microchannel versus 3 sets of sub-100-μm wide microchannels. To do so, we plotted the number of axons exiting the corresponding cluster of microchannels at the two chosen experimental time points. Our results show that at DIV 2, with no extra medium added in the culture chamber (0 μl hydrostatic condition), there are no significant differences in axonal number between the various microchannel widths (Figure 3A). However, the increase of microchannel width leads to a significantly higher number of projected axons at DIV 9 (Figure 3B). These data suggest that, indeed, the presented microfluidic designs have an impact on the density of axons passing through the microchannels.

**Figure 3:**
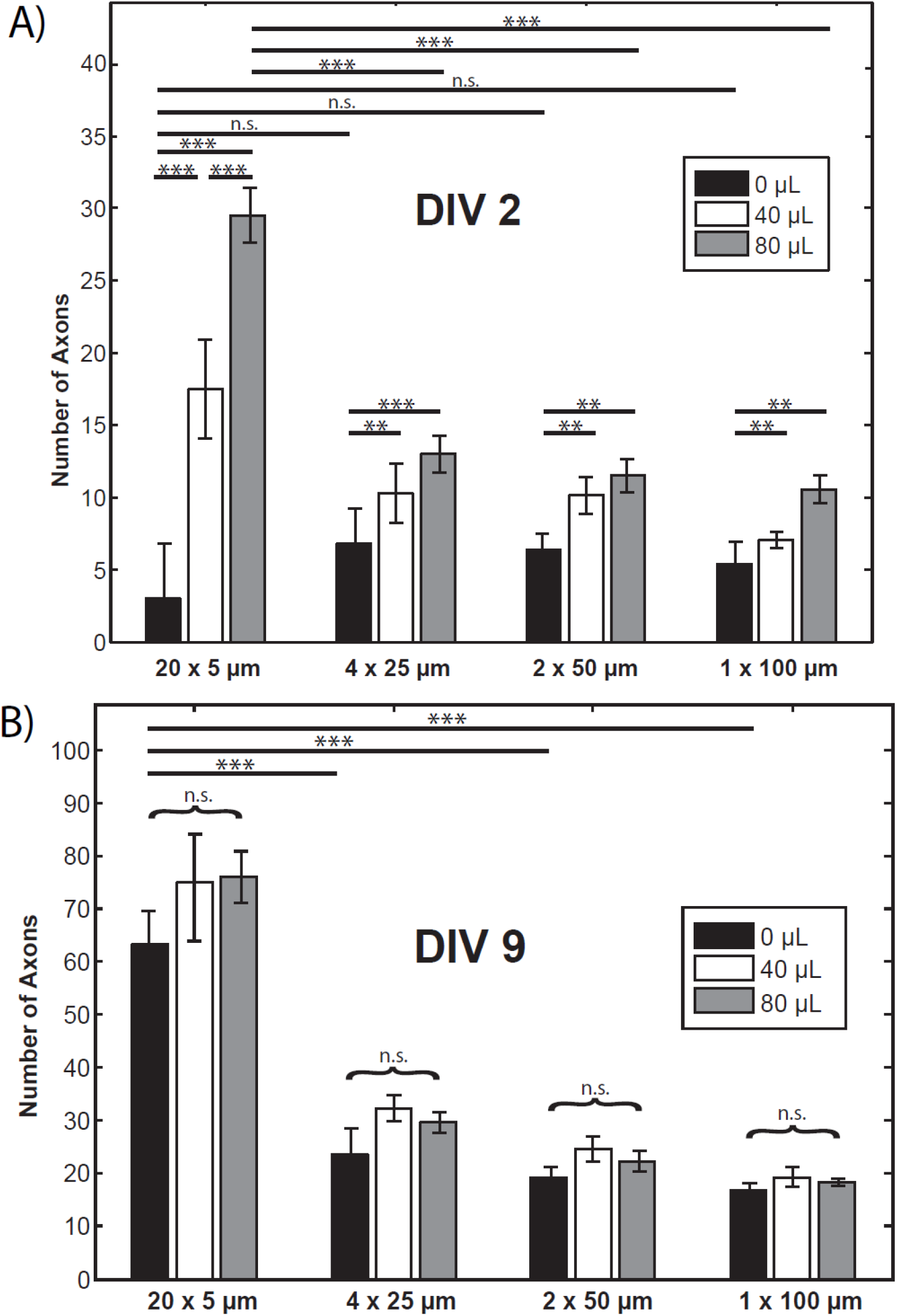
Plots representing the number of axons projected through several sets of microchannels, ranging from 5 to 100-μm wide, according to the three tested hydrostatic conditions at DIV 2 ***(A)***, and at DIV 9 ***(B)***. All data indicated as mean ± SEM. Significance calculated by a post-hoc Tukey’s test; “n.s.” means “not significant”; **p<0.01; ***p<0.001; (n = 3).

It is important to note that, even if the cumulative width is the same along the four different microchannel groups, the amount of projecting axons is significantly higher when comparing 20×5 μm-wide with 4×25 to 1×100 μm-wide microchannel sets (Figure 3B), indicating that axons seem to grow in higher number when passing through clusters of more microchannels of smaller widths. Recent studies have shown that axons prefer to grow alongside physical walls, which is used to fabricate efficient directional axon guides^5,6^. In this regard, it is possible that narrower microchannels increase the physical confinement of developing axons, therefore promoting their unidirectional growth. The growth cones might be able to sense their physical environment more rapidly, allowing them to advance faster through the microchannel. However, this difference in axonal number is lost between sets of wider microchannels (> 25 μm-wide), being the quantity of projected axons stable across the different tested widths (Figure 3B). To explain this observation, we hypothesized that developing axons within a “large” microchannel might be able to grow into alternative directions, allowing them to turn around and exit in the culture chamber of origin. Remarkably, other studies have also described such axonal fasciculations and projections growing in bundles when penetrating wider microchannels, leading to a reduced number of axons exiting on the second chamber^5,6^.

We also investigated the influence of hydrostatic pressure on the axonal projections within the different microchannel widths of our device. At DIV 2, we can observe that, the higher the applied pressure gradient, the more axons are counted in each microchannel exit (Figure 3A, Suppl. Figure 1). These results indicate that axons grow faster when helped with a small pressure gradient, subsequently reaching the second microfluidic chamber sooner. However, this trend is not observed when neurons reach DIV 9 (Figure 3B, Suppl. Figure 1), suggesting that the applied pressure gradients might not influence the final number of axons going through the microchannels. Thus, the axons seem to grow faster and further under a slight hydrodynamic pressure, but this pressure is not enough to increase their number.

### Engineering the structural connectivity pattern of in vitro neuronal cultures

The resulting influence of the different microchannel widths on the density of projected axons in DIV 9 neurons is summarized in Figure 4. In this figure one can observe that, while applying hydrostatic pressure does not seem to change the number of axons of passage, the use of sets of multiple sub-25-μm wide microchannels significantly increases the quantity of projected axons from one culture chamber to the other. In the light of these data, it is possible to effectively engineer the configuration of the microchannels linking two microfluidic chambers in order to maintain a controlled structural connectivity between two neuronal populations. Knowing how to accurately optimize the architecture of this connectivity through adjusting axonal density allows for more flexibility when designing microfluidic devices to construct *in vitro* neural networks.

**Figure 4:**
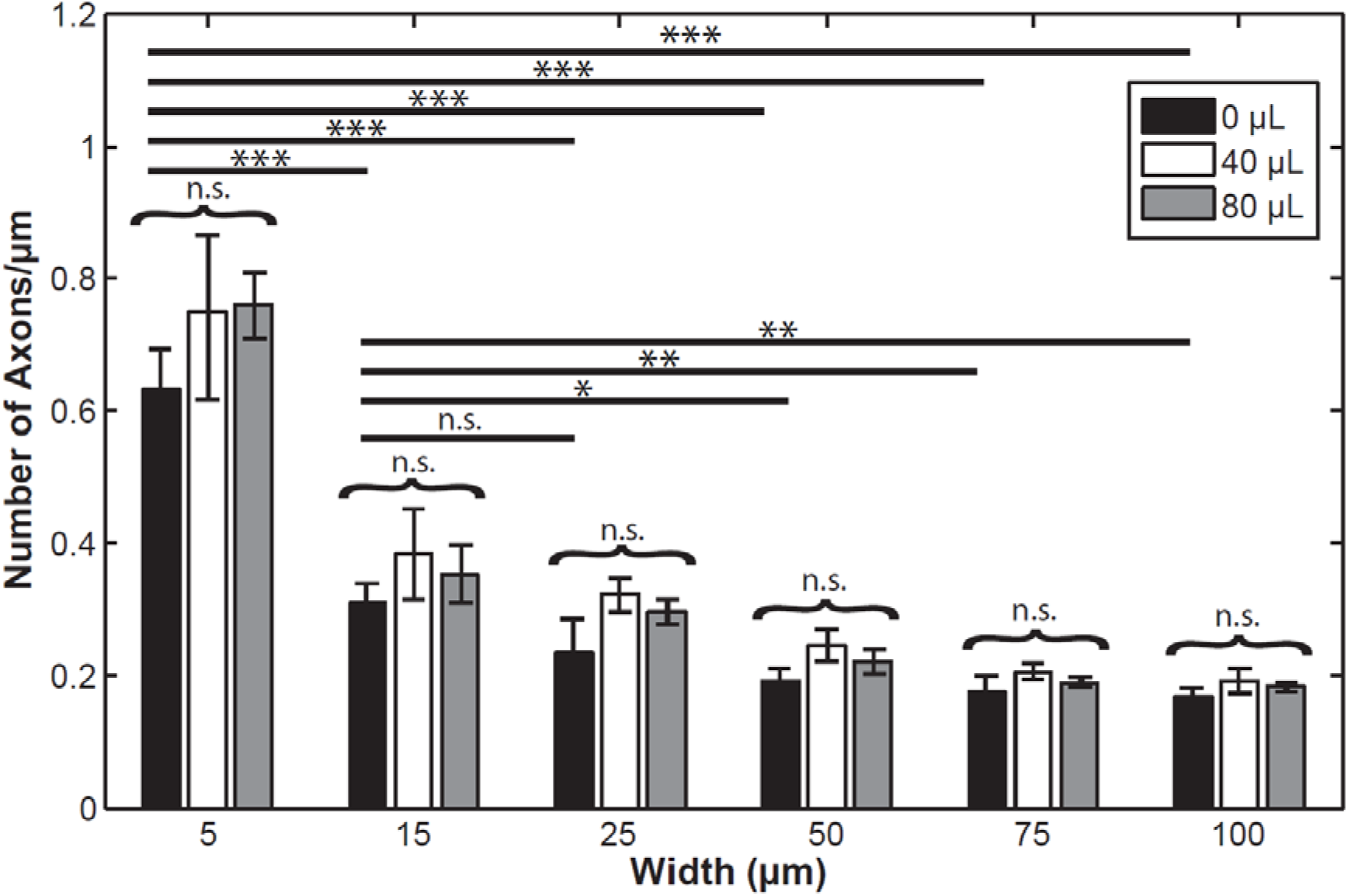
Graph plotting the quantified number of axons projected through all the tested microchannel widths according to the three tested hydrostatic conditions (0 μL, 40 μL, and 80 μL of extra medium added) at DIV 9. All data indicated as mean ± SEM. Significance calculated by a post-hoc Tukey’s test; “n.s.” means “not significant”; *p<0.05; **p<0.01; ***p<0.001; (n = 3).

Being the axonal weight connecting different neuronal populations a key feature of the nervous system organization, our work brings a cutting-edge change in current microfluidic platforms for neurobiological research. Reconstructing a reliable interconnected and oriented neuronal network is an important step for the creation of minimalistic and complex models of the brain^5^. This will not only bridge the gap between *in vivo* and *in vitro* studies, but also will help to understand the structural-functional relationship of neural connections that will enable the high-throughput modelling of neurodegenerative diseases^4^.

To further strengthen the applicability of those strategies, further work should analytically examine co-cultures of neurons and glia^12,17^ and also co-cultures of different neuronal types^5,18^, to account for the cellular heterogeneity present in the nervous system. Likewise, the effect of axonal density variations on the connected neural populations must be verified through monitoring cellular activity using calcium imaging^9,16^ or electrophysiology assays^7,8^.

## Conclusion

In this study, we managed to regulate the number of projecting axons through sets of microchannels with differing dimensional characteristics, therefore defining microchannel patterning design rules to improve current microfluidic approaches for the *in vitro* monitoring of neuronal connectivity. We believe that those strategies will allow the manipulation of the structural connectivity pattern between two or more populations of neurons, and so to examine the effects of axonal density on the neural network dynamics.

With the growing interest from the scientific community to create *in vitro* minimalistic and physiologically relevant neural circuits, the need to control the structural nature of the connections between neuronal populations has never been more pressing. Our work brings a novel method to effectively increase axonal growth using a combination of microchannel width and number variations, without the need of utilizing chemical cues.

In conclusion, these data apport useful engineering design rules for the future investigation of specific connectivity patterns of neural circuits among populations of neurons from different types, which is key for the *in vitro* study of the complexity and integrity of the brain, and also the pathophysiology and treatment of neurodegenerative diseases.

## Acknowledgements

We thank Dr. Clara Berenguer Escuder (CBE Science Writing) for organizing the structure of the manuscript, and for the revision and editing of the original draft. We also thank Louise Miny for her help on editing the article.

