## Supplementary Figure 1 for "Microchannel patterning strategies for *in vitro* structural connectivity modulation of neural networks"

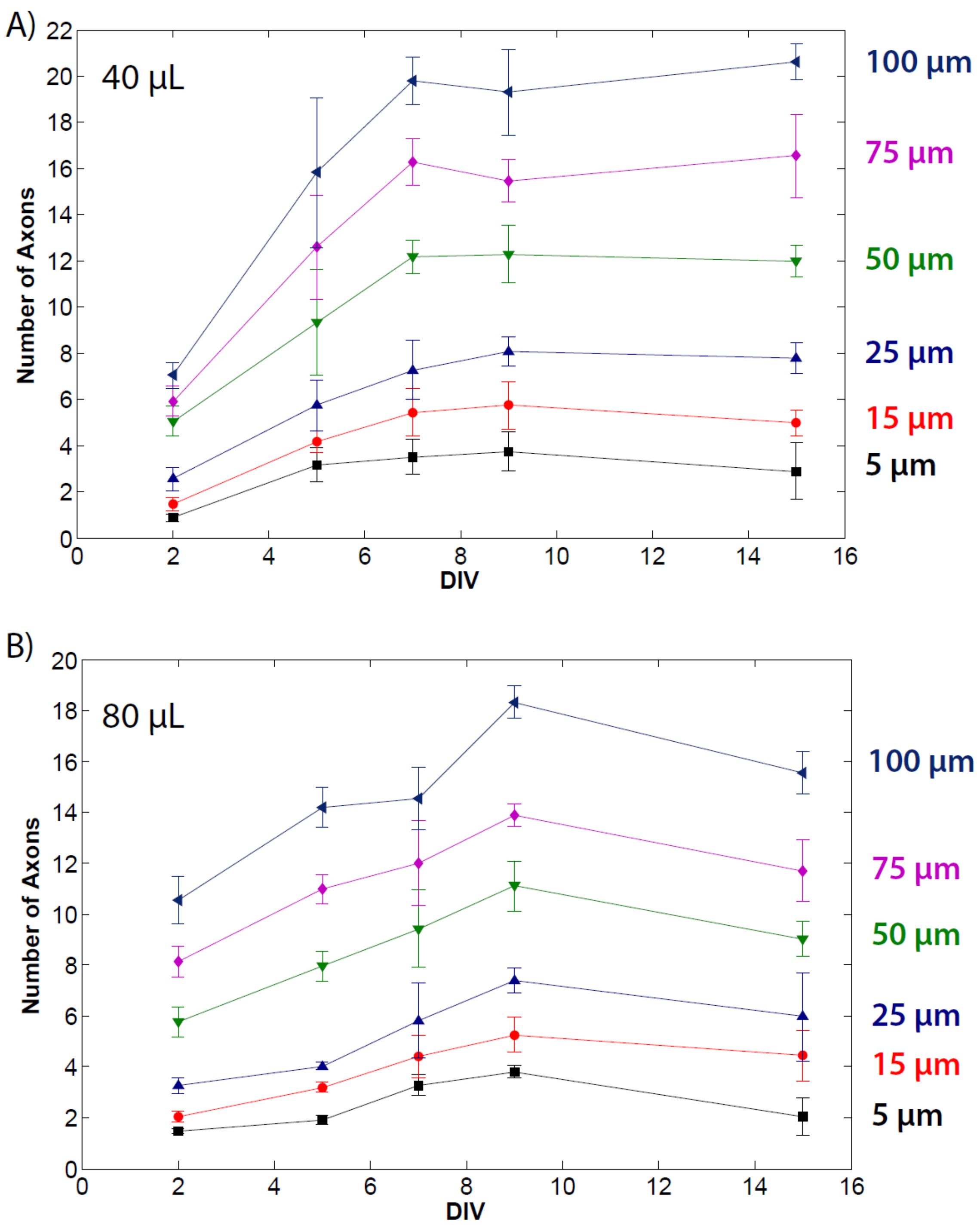


**Supplementary Figure 1:** Quantification of the number of projected axons through the microchannels of various widths as a function of days *in vitro* (DIV) when applying 40 µL ***(A)*** and 80 µL ***(B)*** of hydrostatic pressure. All data indicated as mean ± SEM; (n = 3).
